# Increased parasite load is associated with reduced metabolic rates and escape responsiveness in pumpkinseed sunfish host

**DOI:** 10.1101/2022.01.24.477519

**Authors:** Joëlle Guitard, Emmanuelle Chrétien, Jérémy De Bonville, Dominique G. Roche, Daniel Boisclair, Sandra A. Binning

## Abstract

Wild animals have parasites that can compromise their physiological and/or behavioural performance. Yet, the extent to which parasite load is related to intraspecific variation in performance traits within wild populations remains relatively unexplored. We used pumpkinseed sunfish (*Lepomis gibbosus*) and their endoparasites as a model system to explore the effects of infection load on host aerobic metabolism and escape performance. Metabolic traits (standard and maximum metabolic rates, aerobic scope) and fast-start escape responses following a simulated aerial attack by a predator (responsiveness, response latency, and escape distance) were measured in fish from across a gradient of visible (i.e. trematodes causing black spot disease counted on fish surfaces) and non-visible (i.e. cestodes in fish abdominal cavity counted post-mortem) endoparasite infection. We found that a higher infection load of non-visible endoparasites was related to lower standard and maximum metabolic rates, but not aerobic scope in fish. Non-visible endoparasite infection load was also related to decreased responsiveness of the host to a simulated aerial attack. Visible endoparasites were not related to changes in metabolic traits nor fast-start escape responses. Our results suggest that infection with parasites that are inconspicuous to researchers can result in intraspecific variation in physiological and behavioral performance in wild populations, highlighting the need to more explicitly acknowledge and account for the role played by natural infections in studies of wild animal performance.

## Introduction

Experimental biologists studying wild animals often assume that their subjects are healthy and performing to the best of their abilities (e.g., Wilson et al., 2015). However, at any given moment, animals are host to a range of parasites or pathogens that can compromise their physiological and behavioural performance with significant ecological repercussions (Poulin et al., 1994; Marcogliese, 2004; McElroy and de Buron, 2014; Binning et al., 2017 Timi and Poulin, 2020). For example, infection by the protozoan, *Ophryocystis elektroscirrha*, causes 14% shorter flight durations and 19% shorter flight distances in Monarch butterflies, *Danaus plexippus*, impairing their ability to successfully migrate (Bradley and Altizer, 2005). The problem of infection is not unique to animals caught in the wild. The microsporidium, *Pseudoloma neurophilia*, a common infection in laboratory populations of zebrafish, *Danio rerio*, alters zebrafish shoaling behaviour and startle responses (Spagnoli et al., 2015; Spagnoli et al., 2017). As a result, parasites may be an important, yet overlooked, driver of intraspecific trait variation in both wild and laboratory populations.

The pervasiveness of parasites in both terrestrial and aquatic systems has been repeatedly highlighted in the ecological literature (Poulin and Morand, 2000; Kuris et al., 2008; Caballero et al., 2015). Similarly, their physiological and behavioural effects on hosts can be dramatic. For instance, trophically-transmitted parasites can affect host predator-avoidance or risk-taking behaviours to facilitate transmission to their final host (Kuris, 2003; Blake et al., 2006; Parker et al., 2015). In killifish, *Fundulus parvipinnis*, individuals infected with larval trematodes swim to the surface, jerk, and shimmer more often than uninfected fish, rendering them 31 times more susceptible to predation by birds (Lafferty and Morris, 1996). Although parasites generally have a detrimental effect on host performance capacity (i.e. the ability of an organism to carry out an ecologically relevant tasks; McElroy and de Buron, 2014), infection can also impact hosts in counter-intuitive ways. For example, high loads of the muscle-dwelling myxozoan, *Kudoa inornate*, are related to faster burst-swimming speeds and gait transition speeds in spotted seatrout, *Cynoscion nebulosus* (McElroy et al., 2015). Thus, the effects of parasite infection on individual performance capacity can be difficult to predict.

Performance capacity, including aerobic metabolic performance, can determine individual success inactivities such as foraging, locomotion, reproduction and predator avoidance (Bennett, 1980; Arnold, 1983). Aerobic metabolic performance is tightly linked to an organism’s ability to uptake oxygen and can be estimated by measuring an animal’s oxygen consumption rate 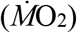 as a proxy of whole-organism metabolic rate (Claireaux and Lefrançois, 2007; Chabot et al., 2016a). Two important physiological traits can be used to describe the upper and lower bounds of an animal’s ability to metabolize oxygen. The maximum metabolic rate (MMR), and the standard metabolic rate (SMR) are defined, respectively, as the maximum and minimum amount of energy that can be metabolized aerobically by an organism (Hulbert and Else, 2000). In ectotherms, SMR is the minimal amount of energy needed for maintenance at a given temperature and is estimated by measuring 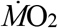 in a non-reproductive, resting, and post-absorptive state (Chabot et al., 2016b). MMR can be estimated by measuring an organism’s 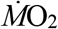 during or shortly after exhaustive exercise (Norin and Clark, 2016; Rummer et al., 2016). The difference between these two traits, the absolute aerobic scope (AS), represents an animal’s ability to perform functions above those required for basic maintenance, including mounting an immune response, digesting, moving, growing and reproducing (Claireaux and Lefrançois, 2007). Parasites that interfere with any aspect of energy demand or physiology might affect the upper and lower bounds of an animal’s AS, and therefore its capacity to carry out various physiological or behavioural tasks. Notably, activation of the immune system during infection may lead to an increase in the host’s SMR, and therefore, reduce its AS (Eraud et al., 2005; Bashir-Tanoli and Tinsley, 2014). Alternatively, parasite infection has also been found to decrease SMR when parasites are located in - or cause tissue damage to - metabolically active organs (Caballero et al., 2015; Ryberg et al., 2020). Similarly, parasites that affect tissues such as the gut, liver or skeletal muscles could impair MMR if they affect the ability of the animal to direct blood flow to these tissues (Coleman, 1993; Gentile and King, 2018). Host anaerobic performance may also be impaired during infection. In response to a predator attack, fishes often perform a sudden burst of anaerobically-powered swimming, known as a fast-start escape response (Domenici and Blake, 1997). Parasitic infection could alter both behavioural (responsiveness, response latency) and kinematic (escape distance, swimming speed, acceleration) components of fast-start escapes. For instance, a recent study reported that experimental infection with a gnathiid isopod ectoparasite in juvenile Ambon damselfish, *Pomacentrus amboinensis*, increased their escape latency to a simulated predator attack by 32% (Allan et al., 2020).

Assessing the effects of parasites on the physiological and behavioural performance of hosts necessitates a proper quantification of parasite load (i.e. the number of parasites in a host, a.k.a. infection intensity). There are many challenges associated with quantifying infection; it is time consuming and often requires detailed knowledge of parasite taxonomy. These reasons may explain why researchers tend to use individual infection status (infected vs. non-infected) rather than their actual loads (i.e. number of parasites in a host), to study the effect of infection on performance. Few studies have explicitly quantified the relationship between host physiological or behavioural performance in wild populations across a gradient of natural parasite infection (but see Ruehle and Poulin, 2019; Ryberg et al., 2020; Sun et al., 2020). Moreover, parasites are often internal, and can only be counted post-mortem. As a result, parasites are not routinely considered in studies on wild animals (Dougherty et al., 2016). This oversight is unfortunate given the high prevalence of parasites in wild populations and their important ecological role (Timi and Poulin, 2020). One means by which some researchers have gotten around this problem is by focusing on the presence of visible infections that are easy to identify. For example, Happel (2019) used photos uploaded to the public database, iNaturalist (www.inaturalist.org), to explore the biogeography of black spot disease in fishes across North America. Black spot disease is caused by infection with the metacercaria of digenean trematodes and can easily be identified and quantified non-invasively through the presence of conspicuous black spots on a fish’s surfaces. Heavy black spot loads in juvenile bluegill sunfish (*Lepomis macrochirus*) causes changes in oxygen consumption rates, body condition, and total body lipid content, reducing overwinter survival to nearly 0% for fish with more than 50 black spots (Lemly and Esch, 1984). These types of infections provide an unparalleled opportunity to consider effects of parasites on wild animals, and thus in experimental research using those animals. Since wild animals are often simultaneously co-infected with several parasite species (Bordes and Morand, 2011), identifying whether “visible” parasite loads, such as black spot disease are related to infection load with other “non-visible” parasites may provide researchers with a simple and useful means of accounting for parasite infections in their studies.

Here, we explored the relationship between parasite infection and aerobic metabolic performance as well as fast-start escape performance in wild-caught, naturally infected pumpkinseed sunfish, *Lepomis gibbosus*. First, we assessed whether visible infections can be used as a proxy of overall endoparasite burden, and thus costs, in hosts, by separately quantifying visible (i.e. trematode metacercaria causing black spot disease) and non-visible (i.e. other cestode and trematode endoparasites) infections in fish. Next, we examined the relationship between parasite load and aerobic metabolism (MMR, SMR, AS) as well as escape performance (responsiveness, response latency, distance travelled) in wild caught fish. Although we were interested in testing for a relationship between visible and non-visible infections, we had no a priori prediction as to the direction of this relationship. Following the overall tendency for parasites to decrease host performance (McElroy and de Buron, 2014), we also predicted that aerobic metabolism and escape performance would be negatively related to greater parasite load in fish hosts.

## Material and methods

### Study species

Sunfishes (*Lepomis* sp.) are abundant in Eastern North America and have been used as model species in behavioural, ecological, and kinematic studies for decades (Brett and Sutherland, 1965; Lemly and Esch, 1984; Tytell and Lauder, 2008; Gerry et al., 2012; Crans et al., 2015). Sunfishes are also hosts to a range of parasites (Margolis and Arthur 1979), which can have a negative impact on whole organism performance capacity (McElroy and de Buron, 2014; Binning et al., 2017). In particular, trematodes causing black spot disease are common in many populations of sunfish (Chapman et al., 2015). The trematodes that cause black spot disease have a complex life cycle requiring two intermediate hosts, typically a snail and a fish, with a piscivorous bird or mammal as the final host (Hunter and Hunter, 1938). Larval trematode cercaria emerge from the snail and encyst under the fish’s skin, in fins and muscle, forming black spots approximately 21 days after infection (Hunter and Hunter, 1938; Hugghins, 1959; Berra and Au, 1978). Pumpkinseed sunfish are hosts to many other endoparasites (e.g., cestodes; including *Proteocephalus sp*, other trematodes including yellow grub; *Clinostomum marginatum*), which can be counted and identified post-mortem (Margolis and Arthur, 1979). Pumpkinseed sunfish naturally infected with black spots provide a great opportunity to assess the degree to which visible (i.e. black spot disease) and non-visible parasites (i.e. other endoparasites) impact the performance capacity of their hosts.

### Fish collection and housing

A total of 42 naturally parasitized pumpkinseed sunfish of similar size (total length: 8.5 ± 0.7 cm; mass 10.24 ± 2.46 g; mean ± standard deviation) were captured with minnow traps and seine nets in Lake Cromwell near the Université de Montréal’s Station de biologie des Laurentides (SBL, Québec, Canada; 45.98898°N, -74.00013°W) in July 2019. Individuals of this size at this location are typically between 2 and 4 years of age (scale-based age determination, unpublished data). Fish were transported to the SBL laboratory facilities within one hour of capture and received a hydrogen peroxide treatment (2.5 ml of 3% H_2_O_2_ *per* litre of freshwater) for 30 minutes to remove ectoparasites, fungus or surface bacteria. Fish were then transferred to a 600L flow-through holding tank (215 × 60 × 60 cm, length × width × height) supplied with water pumped from nearby Lake Croche (45.99003°N, -74.00567°W) and held following a 12 h:12 h light: dark cycle. Water was particle-filtered, oxygenated, and UV-sterilized before entering the holding tanks at a rate of 0.14 to 0.68 m^3^ hr^-1^, allowing a full water replacement every 1 to 4 hours (flow rate adjustments were made to maintain the water temperature near 21°C; actual range:19°C - 21°C). Water temperature and oxygen levels were monitored twice daily (OxyGuard, Handy Polaris, Denmark) and excess food and debris were siphoned daily. Fish were left in the holding tank for 24 h before each individual was measured (wet mass (g), total (TL) and standard (SL) length (mm)). Each fish was identified with a unique three-colour code using visual implant elastomer tags (VIE; Northwest Marine Technology) implanted on each side of the dorsal fin with a 29G needle. Throughout all procedures, fish were manipulated in individual water-filled plastic bags to minimize air exposure and stress. All fish were fed to satiation twice daily (8:30 AM and 6:30 PM) with a mix of bloodworms and commercial fish pellets (Nutrafin Bug Bites, Cichlid Formula) and were habituated for 3 to 5 days before the onset of experiments. After this habituation period, the fish underwent respirometry trials to estimate oxygen consumption. This study was conducted with approval from the Université de Montréal’s animal care committee (Comité de déontologie de l’expérimentation sur les animaux; certificate number 19-034). Respirometry trials Metabolic traits (MMR, SMR, AS) were estimated as rates of oxygen uptake 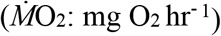 using intermittent flow respirometry. Two identical, separate experimental water baths, (78 cm × 33 cm × 38 cm, length × width × height, 80 L) each contained four resting chambers made of Perspex cylinders (16 × 6 cm, length × diameter). The chambers were opaque with a transparent viewing window located on top. Each chamber was connected to a closed water circuit (491 ml; volume includes recirculation tubes) with a recirculation pump (to achieve adequate water mixing) on which a fiber-optic oxygen probe (firesting 4-channel oxygen meter, PyroScience GmbH, Aschen, Germany) was connected. Dissolved oxygen levels were measured every 3 seconds. The four chambers were connected to a flush pump operated by a digital timer programmed to turn on for four minutes and off for six minutes. This created a 10-minute loop allowing for a four-minute period of water replacement and oxygenation and a six-minute period where the chamber was sealed with no outside exchange of water. A third water bath where temperature was regulated via an aluminium coil pumping chilled water (Thermo Fisher Scientific, EK20 immersion cooler, USA) was used to keep water temperature in the chambers near 21°C (actual range: 20.8°C - 21.7°C). Water replacement in this bath was filed with filtered lake water (same as the holding tanks) pumped through a UV-sterilizer. Background oxygen consumption rates 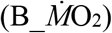 were estimated in each empty chamber for 30 minutes before and after every respirometry trial. The respirometry chambers, tubing, pumps and water baths were cleaned every 3 days with a mix of warm water and 3% hydrogen peroxide (H_2_O_2_) and left to dry outside in direct sunlight.

Fish were fasted for 24 hours prior to all respirometry experiments to ensure they were in a post-absorptive state (Clark et al., 2013; Chabot et al., 2016b). Each trial started with a 3-minute chase protocol followed by 1-minute of air exposure, a common method of estimating MMR in fishes that are poor endurance swimmers (Roche et al., 2013; Rummer et al., 2016). A fish was transferred to a circular chase arena (48 cm × 41 cm, height x diameter, 67 L) using a water-filled plastic bag. The fish was then chased by hand for 3 minutes. When the fish began to fatigue, the experimenter would lightly pinch the fish’s tail to force swimming until it no longer swam. Fish were then removed from the arena and held out of the water for 1-minute. The fish was then placed into a respirometry chamber, which was immediately (less than 15s) sealed for 10 minutes to estimate MMR. Once all eight fish had been chased and the 10-minute measurements completed, control of the system was switched to the automatic timers running the 10-minute loops as described above for the next 18 to 20 hours, during which oxygen consumption rates of fish stabilized, and SMR could be estimated. Oxygen levels remained above 80% in the chambers during all trials. Once a respirometry trial was over, fish were removed from the chambers and returned to their holding tank to recover during 5 days before the escape response trials. This protocol follows best practices for collecting and reporting respirometry data as described in Killen et al., 2021.

### Escape response trials

Escape response experiments were conducted to estimate a fish’s reaction to a simulated aerial predator attack. These experiments were performed between 8:30 AM and 5:00 PM, on fish that had been fasted for 12 to 20 h, to prevent them from regurgitating food during a trial and to maximize the energy available for swimming and recovery. The escape response arena and experimental protocol were based on designs and procedures described in Binning et al., (2014) and Roche, (2021). Briefly, fish were introduced to the escape response arena in a water-filled plastic bag to minimize air exposure. The arena was a 60 × 60 × 30 cm (length × width × height) acrylic clear bottom tank under which a mirror was suspended at a 45° angle to film the escape response from below. The escape response arena was filled with the same water as the holding tanks to a height of 8 cm, which limited vertical movements by the fish while permitting the full extension of their dorsal and pelvic fins. The water temperature in the arena was maintained at 21°C and changed every hour to control temperature and oxygen levels (>95% air saturation). Prior to the experiments, fish were left undisturbed in the arena for 10 minutes to acclimate. We used a mechano-acoustic stimulus located in the far-left corner of the arena to simulate an aerial attack. A weighted stimulus (iron bolt, 2,6 cm long) was released by an electromagnet and fell through an opaque PVC tube (22 cm long and 4 cm wide) suspended 1 cm above the water surface to avoid visual stimulation of the fish (Binning et al., 2014; Marras et al., 2011). Fish were stimulated when they were static (i.e. not swimming), had the stimulus in their field of view and were a maximum of 10 cm from the stimulus. Each individual was subjected to three trials, with a 10-minute interval between trials to allow recovery (see Jornod and Roche, 2015). Escape responses were filmed at 240Hz with a high-speed camera (EX-FH100, Casio, USA). Fish were euthanized following the escape response experiments with an overdose of eugenol solution and placed in a freezer at -18°C until they were dissected.

### Fish dissection

The number of black spots were assessed by counting the number of cysts on the body surface and on all fins visible on the left side of each individual, so that the actual black spot number approximately double of what is reported in this study (Ferguson et al., 2010). Captured fish harboured a varying number of black spots (6-273 metacercaria quantified on the left side of the fish only). Fish body cavity, liver, digestive tract, muscles, and gills were dissected and inspected for internal parasites under a dissecting scope. Internal parasites were counted and identified to the species level. Two species of internal parasites were identified. *Proteocephalus ambloplites* was the most prevalent and abundant species (prevalence: 93%, min-max: 0-153 parasites per fish), and was mostly found in the liver and body cavity. *Clinostomum marginatum* was less prevalent (prevalence: 26%, min-max: 0-7 parasites per fish) and found mostly encysted in the gills and muscle. To correct a fish’s mass for the number of internal parasites it harboured, we weighed approximately 20 individuals of each species of internal parasite and then divided this mass by the number of parasites to obtain an estimate of the mass of one individual parasite. This process was repeated five times with different internal parasites. We averaged these five estimates for each internal parasite species to get a mean individual parasite mass, and then corrected the mass of each fish by the number of parasites of each type it contained (parasite-corrected fish mass; hereafter fish body mass; Lagrue and Poulin, 2015). Metacercaria causing black spots were not weighted as their collective mass was too small to be accurately estimated (±0.000001 g), and likely had no influence on overall fish mass.

### Data extraction and analyses

#### Respirometry data

All oxygen consumption rates 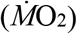 were extracted using the package *respR* (Harianto et al., 2019) in R v. 3.6.1 (R Foundation for Statistical Computing, 2019). Metabolic rate estimates 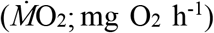 were calculated from the slopes obtained from the linear regression between oxygen concentration and time, accounting for the volume of the respirometer subtracting fish volumes (assuming a density of 1g/ml). The background respiration rate 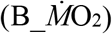 was subtracted from the 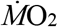 measurements assuming a linear increase in bacterial respiration from the start to the end of the trial. SMR was estimated from measurements taken ∼10h after the onset of the trial (moment at which 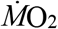 stabilized to a minimum level) until sunrise. The lowest 0.2 quantile of a minimum of 29 slopes (max number = 59) were used to estimate SMR with the *fishMO2* package (Chabot 2016; Chabot et al., 2016b); the mean R^2^ of slopes for all fish was 0.99. We used the *respR* package (Harianto et al., 2019) to estimate MMR with a rolling regression that determines the highest rate of change in oxygen over 60 seconds in the 10-minute measurement following the chase and air exposure protocol, after excluding the first 30 seconds. Absolute AS was calculated as the difference between MMR and SMR (Halsey et al., 2018). Metabolic rates were estimated for 39 fish; data from 3 individuals were excluded because of irregularities in the 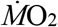 readings due to an air leak.

#### Escape response data

We analyzed the behavioural components of escape responses using VLC media player (VideoLAN, Paris, France). Responsiveness was assessed over the three trials: for each trial, we recorded whether a fish responded to the stimulus (i.e. performed a c-start following contact of the stimulus with the water). Escape latency (sec) was calculated from the number of frames between the first contact of the stimulus on the water and the first head movement of the fish initiating an escape response. Since fish did not respond to the stimulus in all trials, we assessed response latency by recording the best performance (shortest time to respond) of an individual across the three trials (Domenici, 2010). Best performance for the distance travelled (*D*esc) was also used for the analysis.

Stage 1 of the fast-start response started at the first head movement of the fish followed by stage 2 which was defined as the first reversal movement of the head and ended once the fish’s body straightened during the contralateral contraction resulting in a half tail beat (Domenici and Blake, 1997; Eaton et al., 2001). Lolitrack 5 (Loligo Systems, Copenhagen, Denmark) was used to track a fish’s center of mass (CoM) and extract the three following variables: (1) escape distance (*D*_esc):_ distance covered in a fixed amount of time, (2) maximum speed (*U*_max_), and (3) maximum acceleration (*A*_max_). Following the onset of stage one, all variables were measured over 54 milliseconds, which corresponds to the mean duration of stages 1 and 2 for all fish. We used ImageJ (National institutes of Health, Maryland, USA) to estimate (4) a fish’s distance to the stimulus prior to the stimulus touching the water (i.e. the straight-line distance between the fish’s CoM and the center of the stimulus), and (5) the fish’s orientation relative to the stimulus (i.e. the angle formed by a) the linear segment relating the fish’s CoM to the center of the stimulus and b) the linear segment relating the fish’s CoM to its snout (Jornod and Roche 2015). These measures (4 and 5) were included in the models to verify whether the fish’s position relative to the stimulus influenced escape responsiveness, latency or *D*_esc_. We did not examine the effect of parasites on maximum speed and acceleration to reduce the number of statistical tests and since they are the first and second derivative of distance which is analysed in this study.

#### Statistical analyses

All data were analyzed in R v. 3.6.1 (R Foundation for Statistical Computing 2019). Pearson’s correlation coefficient was used to test for a relationship between visible and non-visible infections in pumpkinseed sunfish. We included 44 fish collected at the same time and with the same collection methods from another study. General linear models (LM; lm function in R) were used to model the effect of parasite load on metabolic traits (MMR, SMR and AS). The number of internal parasites, number of black spots, fish body mass (parasite-corrected fish mass), the interaction between the number of internal parasites and fish body mass, and the interaction between the number of black spots and fish body mass were included as predictors in all three models of metabolic traits. Collinearity between fixed factors in the models was assessed using the variance inflation factor (VIF; vif function in *car* package; Fox and Weisberg, 2011). The number of black spots and fish body mass were correlated (n=42; Pearson’s correlation r=0.38, P=0.01), but the VIF terms for these predictors were low (2 at most), so we kept both in all of our models (Legendre and Legendre, 2012). Since fish body mass and fish total length were highly correlated (n=42; Pearson’s correlation r=0.96, P<0.001; Fig. S1), only one of the two was used as a predictor in each model; fish body mass was used in models with metabolic traits as response variables, and total length was used in models with measures of escape performance as response variables.

A general linear model was used to quantify the effect of parasite load on response latency. Response latency was log10 transformed to meet model assumptions. The number of internal parasites, number of black spots, fish total length, distance, and angle of the fish relative to the stimulus, the interaction between the number of internal parasites and fish total length as well as the interaction between the number of black spots and fish total length were included as fixed effects in the model.

A general linear model was used to quantify the effect of parasite load on *D*_esc_. The number of internal parasites, number of black spots, fish total length, distance, and angle of the fish relative to the stimulus, the interaction between the number of internal parasites and fish total length as well as the interaction between the number of black spots and fish total length were included as fixed effects in the model.

We used a generalized linear mixed-effects model (GLMM) with a binomial error distribution (logit link) using the package *lme4* (Bates et al., 2014) to quantify the effect of parasite load on fish responsiveness during escape response experiments. Fish ID was included as a random effect. The number of internal parasites, number of black spots, fish total length, distance, and angle of the fish relative to the stimulus, and the interaction between the number of internal parasites and the number of black spots were included as fixed effects. Covariates in all models were z-transformed using the scale function in R. The angle of the fish relative to the stimulus was sine transformed following Roche (2021). Model assumptions were assessed visually with diagnostic plots and were met for all models (we used functions in the package *DHARMa* for GLMMs; Hartig, 2022): the residuals of all models were normal; no relationship was observed between the residuals and the observed variable and no deviation from the 1:1 line in qq-plots.

## Results

The number of black spots found on a fish’s left side ranged from 6 to 273 (median: 56.5), and the number of internal parasites between 0 and 153 (median: 15). Internal parasite counts include *P. ambloplites* and *C. marginatum* (see fish dissection section). We found no relationship between the number of internal parasites and the number of black spots present in a fish (n=86, Pearson’s correlation r=0.12, P=0.24; Fig. S2). Therefore, visible, and non-visible loads were treated as separate variables in analyses.

Variation in MMR ranged from 2.3 to 7.5 mg O_2_ h^-1^ while SMR ranged from 0.42 to 2.9 mg O_2_ h^-1^. There was a significant positive relationship between all three metabolic traits estimated and fish body mass (Table 1), and no significant interactions between parasite load (black spot and internal) and fish body mass for any of the metabolic traits estimated (Table 1). None of the three metabolic traits estimated was related to black spot load (Fig. 1A, C, E); however, both MMR and SMR were negatively related to internal parasite load (Fig. 1B, D): fish with a higher number of internal parasites had both a lower MMR and SMR (Table 1). There was no relationship between of internal parasite load on AS (Table 1). Number of internal parasites ranged from 0 to 50 for all fish except for one individual harbouring 153 internal parasites. When this individual was excluded from the analysis, both MMR and SMR were still negatively related to internal parasite load (Table S1 and Fig. S6 A, B). Aerobic scope however decreased with internal parasite load when this individual was excluded (Table S1 and Fig. S6 C).

**Table 1.** Test statistics obtained from linear models (LM) of MMR, SMR and AS as a function of black spots, internal parasites (Internal), fish body mass (Mass), the interactions between black spots and mass (BS*mass), the interaction between internal parasites and mass (Int*mass) in pumpkinseed sunfish from Lake Cromwell (n= 39). Statistically significant results are indicated in **bold**. (See table S1 for test statistics for the relationship between parasite load and metabolic traits for fish with 0 to 50 parasites)

| Response | Predictors | DF | F-value | P-value | R<sup>2</sup> |
| --- | --- | --- | --- | --- | --- |
| **MMR**<br>(mgO<sub>2</sub>h<sup>-1</sup>) | Black spot | 1, 33 | 0.82 | 0.37 | 0.67 |
|  | Internal | 1, 33 | 5.15 | **0.03** |  |
|  | Mass | 1, 33 | 59.20 | **<0.001** |  |
|  | Black spot * mass | 1, 33 | 3.38 | 0.08 |  |
|  | Internal * mass | 1, 33 | 0.38 | 0.54 |  |
| **SMR**<br>(mgO<sub>2</sub>h<sup>-1</sup>) | Black spot | 1, 33 | 0.44 | 0.51 | 0.38 |
|  | Internal | 1, 33 | 7.75 | **0.009** |  |
|  | Mass | 1, 33 | 12.64 | **<0.001** |  |
|  | Black spot * mass | 1, 33 | 0.53 | 0.47 |  |
|  | Internal * mass | 1, 33 | 0.88 | 0.35 |  |
| **AS**<br>(mgO<sub>2</sub>h<sup>-1</sup>) | Black spot | 1, 33 | 0.54 | 0.47 | 0.59 |
|  | Internal | 1, 33 | 2.27 | 0.14 |  |
|  | Mass | 1, 33 | 47.02 | **<0.001** |  |
|  | Black spot * mass | 1, 33 | 2.80 | 0.10 |  |
|  | Internal * mass | 1, 33 | 0.13 | 0.73 |  |

**Figure 1.**
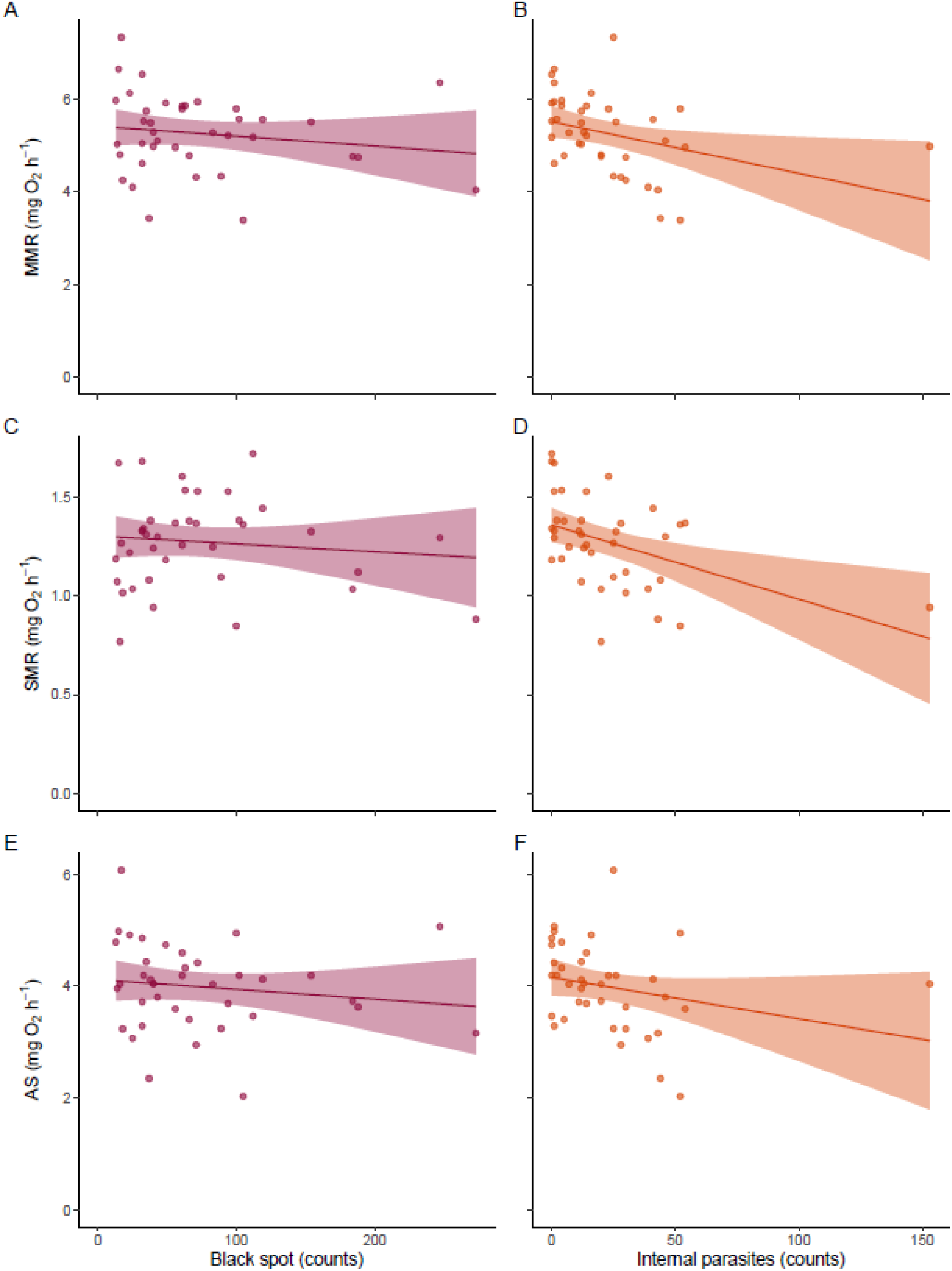
Relationship between host metabolic traits and parasite load. Mass-adjusted metabolic traits (MMR, SMR, AS) as a function of number of black spots (A, C, E) and number of internal parasites (B, D, F) in pumpkinseed sunfish (n=39). Points represent individual fish. The shading around the regression lines represents 95% confidence intervals. Mass-adjusted metabolic rates are metabolic rates (MMR and SMR) adjusted to a common body mass (10.4 g) by adding the residuals of a regression of log MR vs log body mass to the fitted model value for the average body mass of all fish in the study. (See fig. S5 for the relationships between parasite load and metabolic traits for fish with 0 to 50 parasites)

There was no relationship between response latency to an aerial attack and parasite loads (black spot or internal) (LM: n=30, black spot: F=0.13, P=0.72; internal: F=1.05, P=0.31; Fig. 2), length (LM: n=30, F=0.002 P=0.96), or distance and angle of the fish relative to the stimulus (LM: n=30, distance: F=0.07, P=0.79, angle: F=3.73, P=0.07). None of the interactions between blackspots and total length (P=0.09) or internal and total length (P=0.35) were significant.

**Figure 2.**
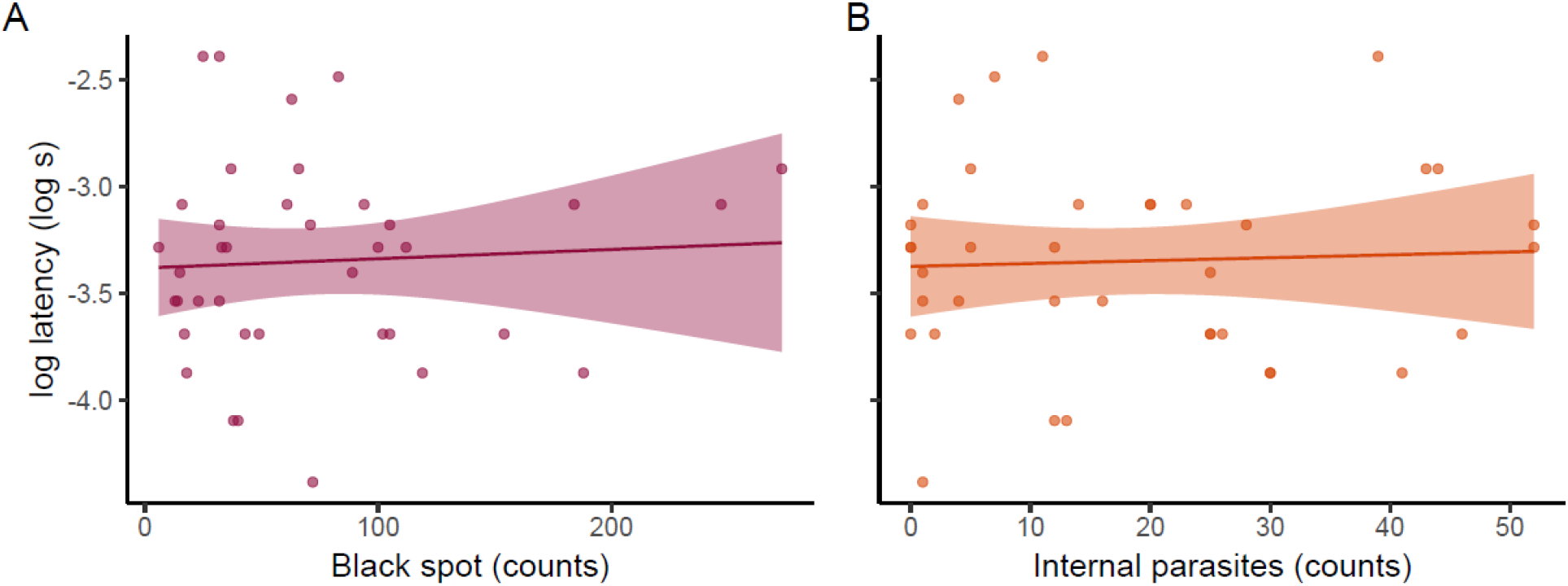
Relationship between response latency and parasite load. Influence of (A) black spots and (B) internal parasites on escape latency in pumpkinseed sunfish from Lake Cromwell (n=38). Points represent individual fish. The shading around the regression lines represent 95% confidence intervals.

There was no significant relationship between D_esc_ and parasite load (black spot or internal) (LM: n=35, black spot: F=1.32, P=0.26; internal: F=0.48, P=0.26), host total length (LM: n=35, F=4.14 P=0.05), distance or angle of the fish relative to the stimulus (LM: n=30, distance: F=0.99, P_dist_=0.33, angle: F=0.26, P=0.61). None of the interactions between blackspots and total length (P=0.21) or internal and total length (P=0.26) were significant.

There were no significant interactions between any of the measured variables in the model with responsiveness to the stimulus as the response variable (Table 2). There was no relationship between fish responsiveness and black spot load (Fig. 3A; Table 2). However, there was a significant negative relationship between fish responsiveness to an aerial attack and fish length (GLMM: n=42, X^2^=10.43, P<0.001; Fig. S3) and a significant effect of the distance of the fish from the stimulus on responsiveness (Table 2). Larger fish responded less often than smaller conspecifics and fish further from the stimulus responded less to the simulated aerial attack. There was also a significant negative relationship between fish responsiveness and the number of internal parasites (GLMM: n=42, X^2^=4.62, P=0.03). Heavily infected fish responded less often to a simulated aerial attack than less infected fish (Fig. 3B). However, this relationship seemed to be driven by two heavily infected individuals (107 and 153 internal parasites respectively). When these individuals were removed from the analyses, this relationship was no longer present (GLMM: n=40, X^2^=0.006, P= 0.94; Fig. 3C).

**Table 2.**
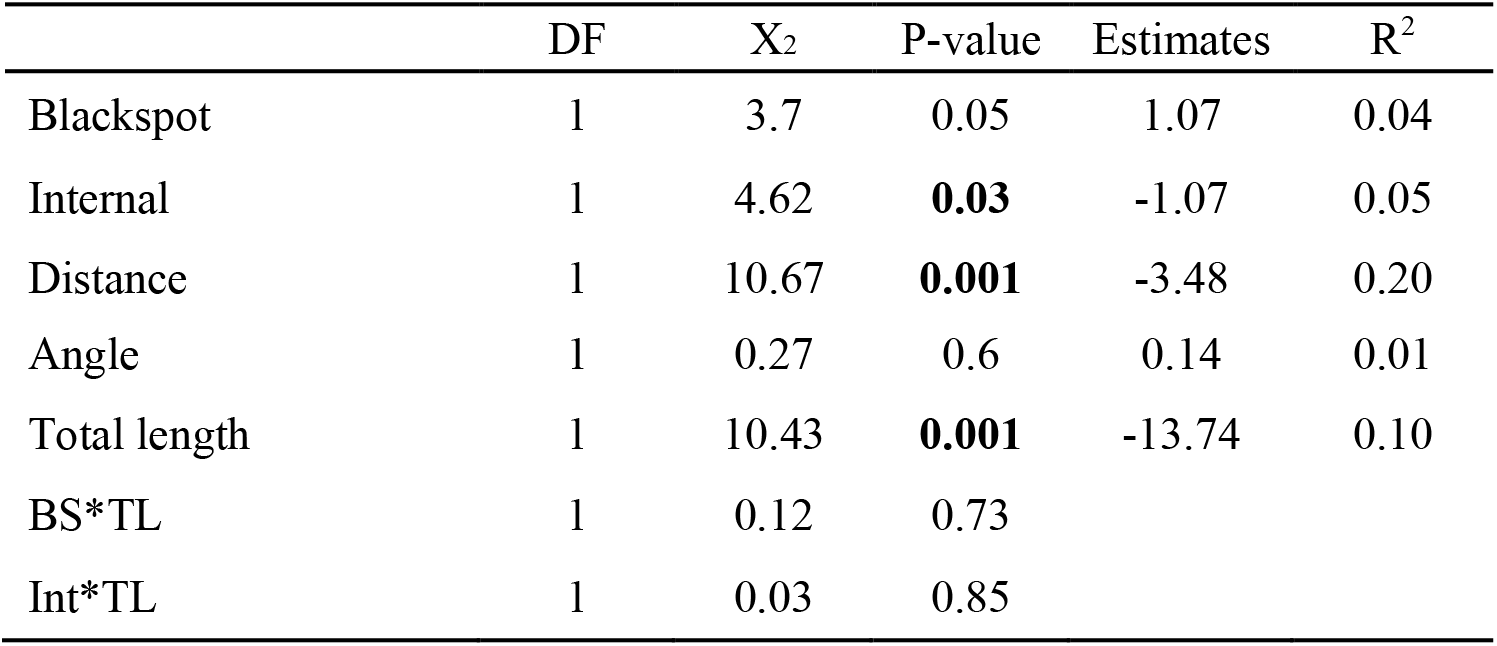
Generalized linear mixed-effects model (GLMM) estimates for black spot load, internal parasite load, distance and angle of the fish from the stimulus and total length (TL) on responsiveness of sunfish from Lake Cromwell. Estimates are from the model without the interactions. Marginal R^2^ for the model = 0.35

**Figure 3.**
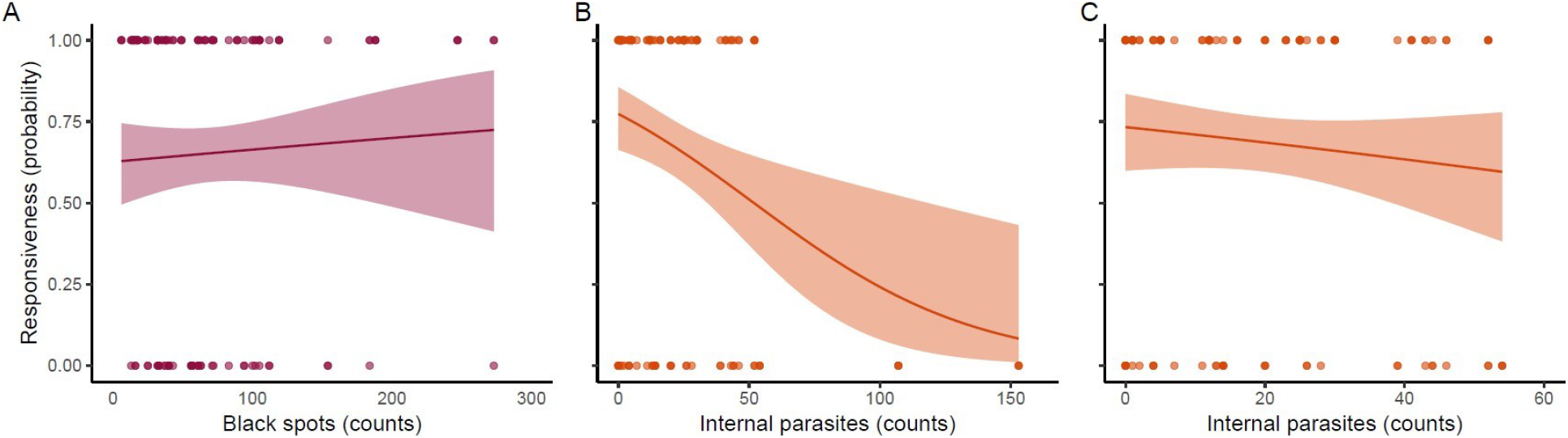
Relationship between responsiveness to a simulated aerial attack and parasite load. (A) Effect of black spot, (B) internal parasites and (C) internal parasites excluding the two most heavily infected individuals on the proportion of trials eliciting a fast-start during escape response experiments in sunfish from Lake Cromwell (n=42). Points represent escape response measurements (up to 3 per fish). The shading around the regression lines represent 95% confidence intervals.

## Discussion

Our results highlight the importance of considering parasite load when studying the physiological and behavioural performance of wild animal populations. We found that metabolic rate estimates such as MMR and SMR, as well as responsiveness to a simulated predator attack decreased along a gradient of non-visible internal parasite infection in pumpkinseed sunfish. This is one of the first studies investigating the impacts of parasite load on aerobic metabolic and escape response performance in adults across two different types of infection (i.e. externally visible black spot infection vs. non-visible internal infections). Our results suggest that experimental studies interested in animal performance may be missing an important driver of intraspecific trait variation by not taking natural parasite infections into account.

### Aerobic metabolic performance

Aerobic metabolic performance traits measured in pumpkinseed sunfish were not related to black spot load. Other studies have also found no noticeable effect of black spot infection on various aspects of host performance capacity. For example, black spot infection did not impact the critical thermal limit nor the body condition of three cyprinid species (Hockett and Mundahl, 1989). Similarily, Vaughans and Coble (1975) found no effect of black spot number on the length-mass relationship, temperature tolerance, or susceptibility of yellow perch, *Perca flavescens*, to predation. Black spot formation is the result of the host’s immune system responding to trematode metacercaria encysting in the host tissues, usually the muscle or fins (Berra and Au, 1978). However, once encysted, metacercaria have very low metabolic costs, and, thus, are unlikely to have long-lasting direct effects on the host’s metabolic traits once the infection is visible (Lemly and Esch, 1984). However, Lemly and Esch (1984) found that the oxygen consumption rates of bluegill sunfish increased approximately one month following experimental infection with the black spot-causing trematode, *Uvulifer ambloplitis*. This corresponds to the average development time (21 days) of the parasite in this host and reflects the time-period during which the parasite likely extracts an energetic toll. Oxygen consumption rates returned to pre-infection levels two months after experimental infection (i.e. one month after the formation of visible cysts) suggesting that the metabolic costs of these infections are short-lived (Lemly and Esch, 1984). A recent meta-analysis also found that the host stress response to parasites is higher early in an infection (O’Dwyer et al., 2020). One rationale for exploring the effects of black spot trematodes on fish performance capacity was to assess whether this visible infection could be used as a proxy for overall infection load and costs given the ease with which black spots can be counted on their hosts. Unfortunately, we did not find a strong relationship between these visible infections and internal parasite load (Fig. 2A) meaning there is no shortcut to quantifying overall parasite load when assessing the impact of infection on individual performance traits.

Although we found no trend between black spot trematodes and aerobic metabolic traits, there were strong negative relationships between the intensity of internal parasite load and metabolic rates. Indeed, we found that both MMR and SMR were lower when internal parasite load was high. This trend was probably driven by infection with *P. ambloplites*, which was much more prevalent and abundant in our population than *C. marginatum* (93% vs. 26% prevalence; 0-153 vs. 0-7 parasites per fish respectively) and more likely to cause extensive tissue damage. *Proteocephalus ambloplitis* tapeworms infect sunfish through the ingestion of infected crustaceans (first intermediate host) such as copepods and cladocerans (Bangham, 1927). The cestode larvae then make their way through the fish’s intestinal walls to the body cavity where they derive energy and nutrients from host tissues including the liver and gonads (Daly Sr. et al., 2006). As such, *P. ambloplites* can cause substantial damage to the organs of its intermediate and final hosts (piscivorous fishes) (Esch and Huffines, 1973; Mitchell et al., 1983). Conversely, *C. marginatum* encysts in the fish’s skin, gills and muscle, and can cause physical damage at the site of encystment due to its relatively large size (3 to 8 mm) (Lane and Morris, 2000). Although the taxonomy and distribution of *Clinostomum* and *Proteocephalus* species has been relatively well studied (Osborn, 1911; Mackie et al., 1983; Muzzall and Peebles, 1998; Caffara et al., 2011; Zimik et al., 2019), little is known about their effects on any of their host’s physiology or behaviour. Our study is among the first to document decreases in physiological and behavioural performance in fishes with high loads of these parasites, which is surprising given their high prevalence and widespread distribution throughout North America.

Standard metabolic rate represents the minimum rate of energy expenditure required to sustain life and sets the floor for an animal’s aerobic metabolic performance (Chabot et al., 2016b). Our results show that parasite infection can be associated with reductions in SMR. Although some studies suggest that parasites tend to increase host energy demands through immune stimulation and maintenance costs (Hvas and Bui, 2022), infection can conversely lead to metabolic suppression in hosts either through a reduction in organ or tissue (e.g. muscle) mass or a decrease in the function of organs associated with energy metabolism (Santoro et al., 2013; Mehrdana et al., 2014; Ryberg et al., 2020). Although a lower SMR can be advantageous in scenarios where food or oxygen are limited (Killen et al., 2016), reduced SMR associated with high parasite loads is more likely a pathological consequence of infection; since much of an individual’s SMR is used to maintain internal organ function, damage caused by parasites can reduce organ function and thus, SMR (Hulbert and Else, 2000; Seppänen et al., 2008; Behrens et al., 2014;; Ryberg et al., 2020). For example, Eastern Baltic cod, *Gadus morhua* infected with high intensities of the liver nematode, *Contracaecum osculatum*, displayed lower SMR, reduced albumin to globulin ratio and lipid content suggesting that the metabolic function of this organ is compromised by high parasite loads (Ryberg et al., 2020). Similarly, *P. ambloplites* were mostly found in our fish’s liver, which was often damaged when infection loads were high. This suggests a direct effect of *P. ambloplites* infection on host aerobic metabolic performance in these sunfish. Experimental infections are needed to establish a causal link between infection and decreased host performance in this system.

Maximum metabolic rate sets the ceiling for aerobic metabolic performance and is associated with increased performance during energetically demanding activities and in high-energy environments (Eliason et al., 2011; Binning et al., 2014; Norin and Clark, 2016). Our results show that internal organ damage caused by endoparasites likely reduces both MMR and SMR in hosts. Studies across taxa report decreases in host MMR with parasite infection (e.g. Careau et al., 2012; Bruneaux et al., 2017; Hvas et al., 2017). Importantly, a decrease in MMR is also often associated with a decrease in AS (Norin and Clark, 2016). We did not observe a decrease in AS with increasing internal parasite load across the entire range of internal parasites recorded (0 to 153 internal parasites), probably due to the lower MMR and SMR observed in heavily infected fish. This suggests that the lower MMR with increasing parasite load decreases somewhat faster than that of SMR, although this result remains marginal. Nevertheless, removing the most infected fish still resulted in an observable negative relationship between metabolic rates (MMR, SMR) and internal parasite load over a range of 0 to 50 internal parasites. Reduced AS can result in less capacity for growth, reproduction and, potentially, survival of heavily infected individuals (Metcalfe et al., 2016). These relationships, and the potential ecological consequences in parasitized individuals, need to be explored more thoroughly.

### Escape behaviour

Responsiveness to a simulated aerial attack was negatively correlated with internal parasite load, but not black spot trematodes. When startled, many fish species perform a characteristic C-start escape response, which is an important determinant of an individual’s survival during a predator attack (Domenici et al., 2011). Escape distance and latency to respond to this attack are all considered important parts of this reaction (Domenici et al., 2011), and can all be impacted by infection (Allan et al., 2020). Yet, we found no relationship between parasite infection and response latency in our adult sunfish. Other studies on adult fish have also found no effect of parasites on escape performance. In bridled monocle bream, *Scolopsis bilineata*, parasitized by Anilocra isopod ectoparasites, response latency, maximum velocity, maximum acceleration and cumulative distance travelled did not differ between infected and uninfected fish (Binning et al., 2014). Similarly, Ruehle and Poulin, (2019) failed to detect a significant reduction in escape performance in infected common bully, *Gobiomorphus cotidianus*, even when host vision was affected.

Although the kinematic components of an animal’s escape response offer useful predictors of an individual’s escape performance, escape responsiveness is arguably the most important determinant of survival in the face of a threat (Domenici, 2010): an individual that does not react to an attacking predator has almost no chance of survival regardless of how fast it can escape. In our study, the two most heavily infected individuals never responded to our simulated aerial attack. Although the negative relationship we observed between responsiveness and parasite load is driven by the escape performance of these two individuals, our results remain ecologically relevant: we collected very few other heavily infected individuals possibly because these individuals are selectively removed from the population through predation. Over-dispersion of parasites within hosts, whereby few individuals harbour most of the parasites in a population, is a well-documented ecological phenomenon (Anderson and Gordon, 1982). Although many factors can explain such patterns, including increased susceptibility and/or tolerance to infection in some individuals, our results suggest that performance reduction in heavily infected individuals may also play a role. If heavily infected individuals are predated upon at higher rates than uninfected or lightly infected individuals due, in part, to decreased responsiveness, we would expect to sample fewer of these individuals in a given population. This phenomenon would also facilitate trophic transmission and therefore be beneficial to the parasite life cycle.

### Other considerations

Host life stage can play a large role in individual responses to parasites. For example, juvenile chipmunks, *Tamias striatus*, show a 7.6% increase in resting metabolic rate in response to infection by botfly larvae, resulting in a ∼5g body mass loss over summer (Careau et al., 2010) whereas no effect of infection is observed in adults (Careau et al., 2012). It is possible that younger individuals are more affected by stressors, including infection, because more energy is required for growth and development (Careau et al., 2010; Allan et al., 2020). Older hosts also typically harbour more parasites than younger ones, probably because parasites are recruited faster than they die in hosts, especially in the case of encysted parasites such as those causing black spot disease (Hawlena et al., 2006). The fact that we did see strong relationships between internal infections and our performance measures, even in our adult fish, is a further reminder of the important impact that parasites can have on their hosts, and their contribution to otherwise unexplained intraspecific variation in performance often observed in natural populations (Timi and Poulin, 2020).

Our study explicitly quantified the load of infection of both externally visible and internally non-visible parasites. However, the process of counting and identifying endoparasites is difficult and time-consuming, and often not included in the context of studies on wild populations. When infection is considered, the host’s infection status (i.e. infected or not) is typically the variable of interest. This binomial categorization can be relevant and related to impact of parasites on some host populations, especially in the case of large endo- or ectoparasites (Fogelman et al., 2009; Jolles et al., 2020). However, parasite load can be more important than infection status for understanding the physiological, behavioural and ecological effects of parasites on their hosts (Poulin, 2019; Timi and Poulin, 2020). For instance, killifish, *Fundulus parvipinnis*, infected with larval trematodes face higher rates of predation by birds along an infection gradient (predation rates in uninfected hosts: 0.02%, lightly infected hosts: 22%, heavily infected hosts: 80%) (Lafferty and Morris, 1996). As there are generally fewer highly parasitized individuals in natural host populations (Crofton, 1971; Shaw et al., 1998), it can be difficult to accurately estimate the effect of high parasite loads on populations because these individuals can be hard to sample and test in the lab. Heavily infected individuals may also be more susceptible to environmental stressors, which could potentially lead to selective mortality based on infection status during extremeevents such as freezing or heat waves (Lemly and Esch, 1984; Bruneaux et al., 2017; Greenspan et al., 2017). The effects of concomitant stressors like temperature and parasite load have rarely been tested in natural populations and remains an area in need of further research especially given projected future global change.

## Conclusion

Our results suggest that parasite load is an important, overlooked driver of intraspecific performance trait differences in host populations. Experimental infections are needed to confirm the causal relationship between infection load and performance traits in fish hosts. We expect experimental infection with black spot and/or internal parasites would result in similar performance trait impairments as documented here. The fact that we were unable to link externally visible black spot infection with non-visible internal parasites is potentially problematic for experimental biologists since these visible infections are a poor proxy of overall infection load. Also, non-visible internal infections, which seem to be related to the highest performance costs, are less likely to be taken into consideration by experimental biologists. While we acknowledge that sacrificing individuals to quantify endoparasite infection is not always feasible or desirable in the context of experimental work on wild animals, we encourage researchers to consider alternative ways of controlling for this potential confounding effect such as treating experimental animals with anti-parasites treatments like praziquantel (Bader, 2017) prior to testing their performance after confirming that such treatments themselves do not impact the traits to be measured.

## Acknowledgements

We thank the staff of the Station de biologie des Laurentides (SBL) de l’Université de Montréal, Gabriel Lanthier, Victoria Thelamon, Isabel Lanthier and Tom Bermingham for field and logistic support; Alexandra Kack, Xue Han Qu, Kaitlin Gallagher and Sean Locke for parasite identification; Amélie Papillon for help with fast-start analysis; and Shaun Killen for helpful comments and advice.

## Data availability

All data presented in this study are publicly available and can be downloaded here: doi: 10.6084/m9.figshare.19005971

